# Welfare concerns for mounted load carrying by working donkeys in Pakistan

**DOI:** 10.1101/2022.03.07.483371

**Authors:** Syed S. U. H. Bukhari, Sarah M. Rosanowski, Alan G. McElligott, Rebecca S. V. Parkes

## Abstract

Working donkeys (*Equus asinus*) are vital to people’s livelihoods. They are essential for carrying goods, however globally, overloading is one of the primary welfare concerns of working donkeys. We studied mounted load carrying by donkeys and associated factors in Pakistan. A cross-sectional study of donkey owners (n = 332) was conducted, and interviews were undertaken based on a questionnaire. Owners estimated that the median weight of their donkeys was 110kg (interquartile range (IQR) 100-120kg), and that they carried a median mounted load of 81.5kg (IQR 63-99kg). We found that 87.4% of donkeys carried a load above 50% of their bodyweight ratio (BWR), the median BWR carried was 77.1% (IQR 54.5-90.7%), and 25.3% of donkeys carried above 90% BWR. Donkeys that were loaded at more than 50% BWR were more likely to sit, compared to donkeys loaded with less weight (p=0.01). Donkeys working in peri-urban and urban areas were more likely to carry a greater BWR than donkeys working in rural areas (P<0.001), as were those carrying construction materials or bricks, compared to agricultural materials (p=0.004). Age (p=0.03) and breed (p=0.01) were also associated with carrying a higher weight. Overloading based on current recommendations (50% BWR) was common, with the majority (87.4%) of donkeys reported to carry more than the recommended 50% limit. This survey provides evidence of on-the-ground working practices and factors associated with mounted load carrying, which is critical for developing evidence-based recommendations for loading, in order to improve the welfare of working donkeys.

## 1 Introduction

Donkeys have played an essential role in developing human civilizations (1). There are approximately 50.5 million donkeys globally (2), benefiting around 600 million people and playing a vital role in the livelihood of poor and vulnerable communities (1,3,4). The importance of working donkeys for their owner’s livelihood and the economies of developing countries is well known (3–5). However, their importance has often been overlooked in government-level animal welfare policies (6). As such little is done to safeguard donkey welfare (7), leading to compromised welfare due to marginalization, harsh working conditions and lack of legislation (3,5).

Donkeys are used in a variety of settings, across rural, peri-urban, and urban areas (7) for the transportation of construction, agricultural, and domestic loads (3–5), including brick production (Figure 1). One of the most severe problems working donkeys experience is overwork and overloading (8–10). Overloading can be defined as the amount of weight that disrupts gait rhythm, resulting in lameness and behavioral changes (5). Donkeys carry tons of weight every day, which likely exceeds their natural weight carrying ability (5), and they work for extended periods of time (3,5,11). Overloading has been identified as impacting on equid behavioral, biochemical, biomechanical, and physiological characteristics (5). High workload and unsafe practices contribute to poor working donkey welfare (3,5).

**Figure 1:**
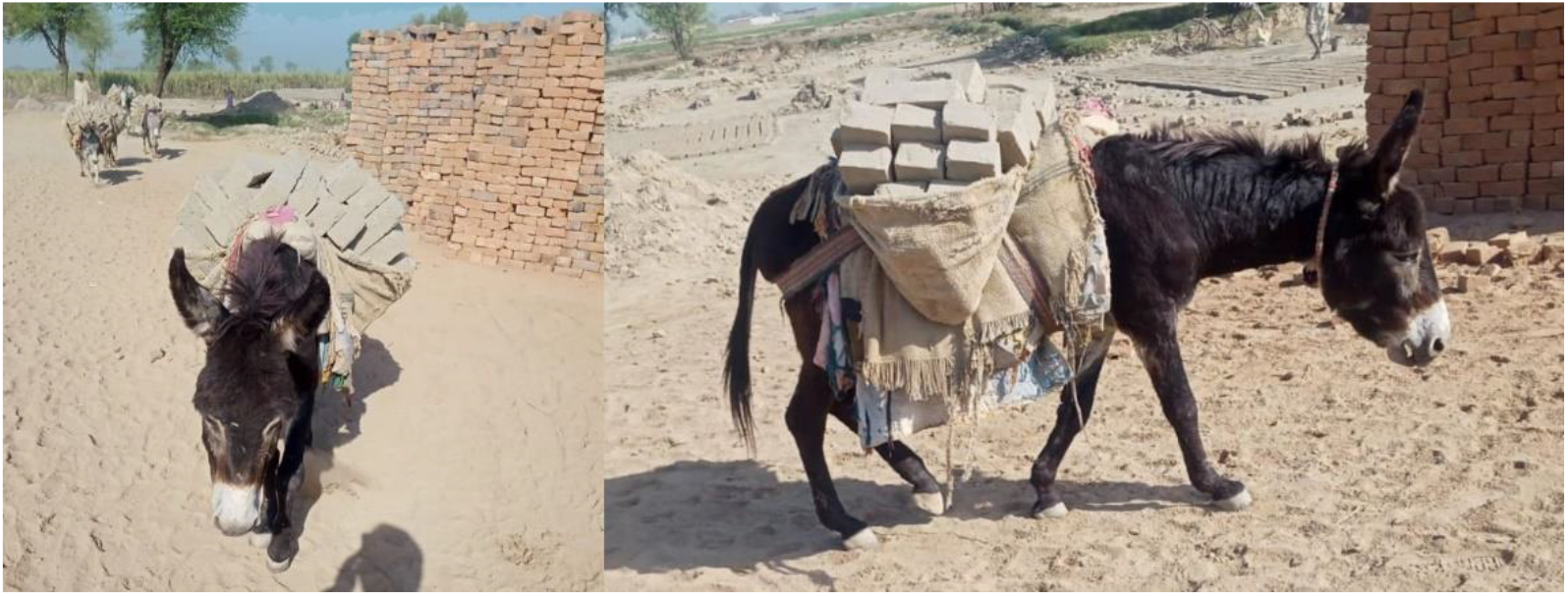
Mounted load-carrying donkeys in a brick kiln production system in Pakistan. Photo: Syed S. U. H. Bukhari

There is little research regarding mounted load-carrying limitations of working donkeys. The maximum load recommended for a fit donkey in the UK is 50kg (12), which is approximately 28% of an adult donkey’s bodyweight, and only when the load is well balanced on its back (12). This 50kg recommendation is not evidence-based, and refers to donkeys in the UK, which typically are in good body condition and are larger (13) than working donkeys in lower-middle income countries (LMICs) (14). Current guidelines for working donkeys suggest that donkeys can safely carry loads of up to 50% bodyweight (15). Donkeys in some LMIC have been reported to carry as much as 75% of their bodyweight (16). However, there is evidence of donkeys carrying up to 117% of their body weight in Pakistan (16,17). Even conservative estimates would indicate that these donkeys carry more than their own bodyweight, which is one of the causes of compromised donkey welfare (5). The current study aimed to quantify demographics of donkey owners, donkey loading practices, and factors related to mounted load carrying in Pakistan.

## 2 Materials and methods

### 2.1 Study area and study design

We carried out a cross-sectional survey of donkey owners in four of Pakistan’s regions (Swat, Attock, Faisalabad, and Bahawalpur; Figure 2). These regions were selected based on their topography and varying climatic conditions: mountainous, arid, irrigated plains, and sandy desert, respectively (18). Different topographic regions were selected because working equids face different challenges in different communities and geographic sites (6,19). The four regions cover almost 39,815km^2^ (almost 4.5% of country) of Pakistan. Swat (34°45’ latitude, 72°54’ longitude) is a mountainous region with an elevation of 2591m above sea level. The maximum average monthly temperature (37 C°) remains during July, and the minimum average monthly temperature (0 C°) is recorded during January. In Swat, annual rainfall ranges between 1200-1400 mm (18). Attock (32°55’ latitude, 72°51’ longitude) is an arid and semi-hilly region with an elevation of 519m above sea level. The maximum average monthly temperature (38 C°) occurs in June, with the minimum average monthly temperature (3 C°) recorded in January. In Attock, annual rainfalls range between 900-1000 mm (18). Faisalabad (31°26’ latitude, 73°08’ longitude) is among the irrigated plains of Pakistan, with an elevation of 185m above sea level. The maximum average monthly temperature (41 C°) occurs in June, with the minimum average monthly temperature (5 C°) in January. In Faisalabad, annual rainfalls range between 300-400 mm (18). Most of Bahawalpur (28°39’ latitude, 70°41’ longitude) is a sandy desert region with an elevation of 88m above sea level. The maximum average monthly temperature (42 C°) occurs in June, with the minimum average monthly temperature (4 C°) in January. In this region, annual rainfalls range between 100-150 mm (18).

**Figure 2:**
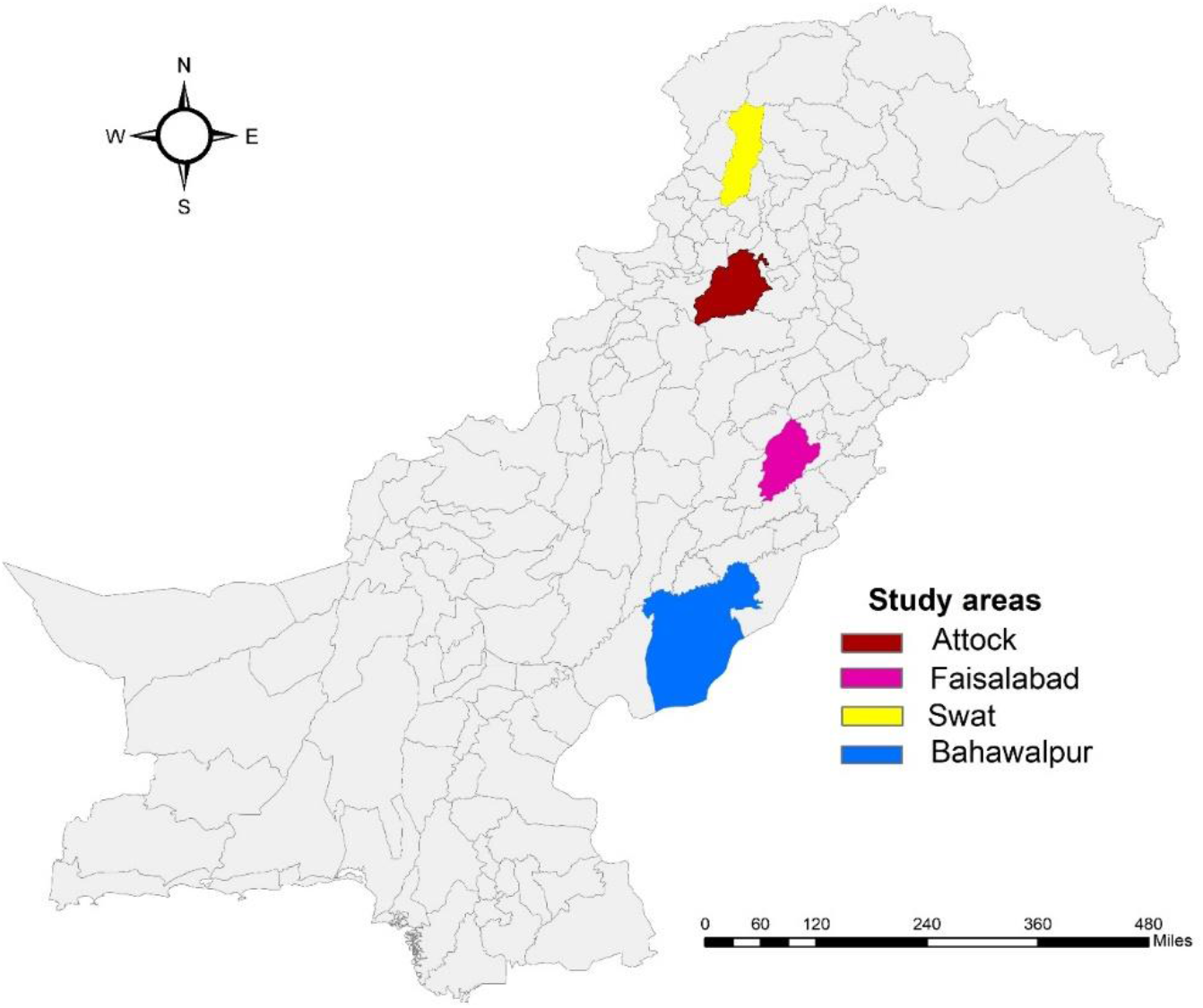
A map of Pakistan showing the locations of the four regions in which the study was conducted.

#### Questionnaire design

A questionnaire was developed to assess demographic characteristics, practices, and factors associated with mounted load carrying. The questions were designed based on field conversations with donkey owners combined with recent field experience of registered equine veterinarians in the four selected study regions. The questionnaire consisted of both open-ended and closed questions. The first section of the questionnaire consisted of informed verbal consent of donkey owners. The second section included information regarding the demographics of the owner, and signalment of the donkey, including type of work undertaken. The third section contained questions on the loading practices. The bodyweight of donkeys and the weight of any mounted loads as a part of their regular loading practices were estimated by the donkey owners. In some cases, donkey owners are able to weigh their donkeys on scales at nearby dairy farms or had recently weighed their donkey. Other owners estimated donkey weight. The weight of the mounted load was estimated depending on the items carried. For bricks, mounted load weight estimation was done by multiplying the number of bricks by the known weight of one brick. The weight of commercially packaged items was written on packaging, for example, a bag of cement, a bag of wheat grain, bags of fertilizers etc. The weight of liquids such as milk, oil, and water containers were estimated by number of liters in one can and the number of cans carried. Donkey owners were asked about two behaviors – whether the owner had previously noted a donkey trying to sit after loading (as sometimes donkeys try to sit by adopting sternal recumbency after loading), or lameness. These two behaviors could be observed by the owner without knowing the cause and to ensure clarity, the behaviors were explained if needed.

A pilot study was conducted, to optimize questions being asked, address discrepancies and check how much time the questionnaire took to complete, by collecting information from 24 randomly selected donkey owners, six from each of the four target regions (20). The time to complete the survey was eight to ten minutes. Surveys were all conducted verbally due to low literacy rates. None of the data gathered from the pilot survey was included and they were not weighed in the final analysis.

### 2.2 Data collection

The survey was conducted by equine veterinarians. They verbally explained the study, its purpose, and its methods. The donkey owners were approached based on convenience sampling and willingness to participate. Once donkey owners had provided consent to participate, interviews were undertaken based on the pre-designed questionnaire. A total of 332 donkey owners participated. They had the opportunity to ask questions, and all their questions were answered appropriately. The interviewer signed a ‘participant informed verbal consent form’. A third person signed the witness statement (witness, to ensure appropriate exchange of information) on ‘participant informed verbal consent form’ according to existing survey guidelines (21,22). Face-to-face interviews were conducted to collect the required information, based on the pre-structured questionnaire which was in English. However, interviews were delivered in the local languages (Urdu, Pashtu, Hindko, Pothwari, Punjabi, Saraiki) after translation by the interviewers who were equine veterinarians and fluent in both English and the respective local languages. This approach was used to maximize the accuracy of responses and minimize any confusion concerning the scientific terminology used according to existing survey guidelines (21,22).

### 2.3 Statistical analysis

The continuous data (weight of donkey, weight of the load, herd size, daily income generated by the donkey, and distance traveled per day) were presented in the form of median, interquartile range (IQR), minimum, and maximum. All the categorical data were described as frequency and percentage.

#### Outcome variables

The following formula was used to calculate the percent bodyweight ratios (%BWR) for all donkeys,

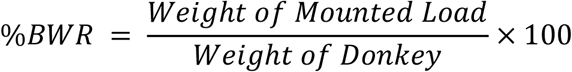

Three new binary outcome variables were created and labeled 1) 50% BWR, 2) median %BWR and 3) high %BWR. Fifty percent BWR was defined as a load of 50% BWR and was selected as an outcome variable based on existing guidelines which suggest that a donkey can safely carry up to 50% of their bodyweight (15). Median %BWR was the median of percent BWR in the population investigated, with half of the donkeys carrying above the median %BWR. The outcome high %BWR was loads at 90% of BWR and represented the upper quartile of our study population.

#### Exposure variables

The variables considered in the logistic regression models were area (urban, peri-urban and rural), donkey age, donkey sex, breed (Sperki, Shinghari, Indian and mixed), type of saddle used, working terrain (mixed, plains, steep) and speed (walking or trotting). Continuous variables were non-normally distributed and were included in the model as categorical variables based on quartiles. Continuous variables were distance covered per day (in km), daily working hours, and earnings of a donkey in Pakistan rupees (PKR). Two donkey behaviors, sitting when loaded (yes/no) and lameness signs while working (yes/no) were included. Donkey breed and age was further categorized as binary variables, i.e., mixed breed and other breeds during multivariable modelling.

#### Univariable and multivariable regression models

Univariable and multivariable logistic regression models were used to determine explanatory variables associated with mounted loads. Three multivariable logistic regression models were developed to investigate factors associated with each of the outcome variables - high %BWR, median %BWR, and 50% BWR. Exposure variables were screened using univariable logistic regression model for each outcome variable. Exposure variables with a likelihood ratio test (LRT) P-value <0.20 were selected for inclusion in the multivariable model for that outcome. A preliminary multivariable model was built using a manual backward stepwise method of elimination in which variables were retained in the final model if the LRT P-value was <0.05. The LRT was used as the primary selection criterion. Confounding was assessed throughout the multivariable model building, with variables changing the odds ratio (OR) more than 10% retained in the final model. The goodness-of-fit of the logistic regression models was assessed using the Hosmer-Lemeshow test. All statistical analyses were conducted using Stata IC version 17 (23).

### 2.4 Ethical approval

This study was approved by the Human Subjects Ethics Sub-Committee, City University of Hong Kong (Approval reference no. JCC2021AY003).

## 3 Results

### 3.1 Demographic characteristics

In total, 332 donkey owners agreed to participate. The demographics of the owners and donkey signalment are presented in Table 1. The majority of questionnaire participants (98.5%; n= 327) were men. Both male (54.5%; n=181) and female (45.2%; n=150) donkeys were used for load-carrying work. The majority of donkeys (58.1%; n=193) were aged between 6 to 10 years. Donkeys worked in rural (48.7%; n=162), peri-urban (38.3%; n=127), and urban (13.0%; n=43) areas. The distance covered by donkeys during their working day was a median of 8km (IQR 3-17km) km. Daily earnings were a median of 685 PKR (IQR 450-900) (USD$3.87 (IQR $2.54 – $5.08).

**Table 1:** Demographic characteristics of the donkey owners and donkeys

| Variable | Level | Number | Percentage (%) |
| --- | --- | --- | --- |
| Owner age (years) | < 31 | 75 | 22.6 |
|  | 31-40 | 141 | 42.5 |
|  | 41-50 | 97 | 29.2 |
|  | > 50 | 19 | 5.7 |
| Owner Gender | Male | 327 | 98.5 |
|  | Female | 5 | 1.5 |
| Area | Rural | 162 | 48.8 |
|  | Peri-Urban | 127 | 38.3 |
|  | Urban | 43 | 13.0 |
| Age of donkey (years) | < 6 | 38 | 11.4 |
|  | 6-10 | 193 | 58.1 |
|  | 11-15 | 62 | 18.7 |
|  | 16-20 | 23 | 6.9 |
|  | > 20 | 16 | 4.8 |
| Donkey sex | Male | 181 | 54.5 |
|  | Female | 150 | 45.2 |
|  | Gelded | 1 | 0.3 |
| Donkey breed | Sperki | 26 | 7.8 |
|  | Shinghari | 49 | 14.8 |
|  | Indian | 9 | 2.7 |
|  | Mixed Breed | 248 | 74.7 |

### 3.2 Mounted loads and %BWR

Median weight for donkeys was 110kg (IQR 100-120kg) and the median mounted load for one trip was 81.5kg (IQR 63-99kg) (Figure 3). The median %BWR was 77.10% (IQR 54.50-90.70%). Overall, 87.4% donkeys carried loads above 50% BWR. Twenty-five percent of donkeys carried loads above 90 %BWR (high %BWR).

**Figure 3:**
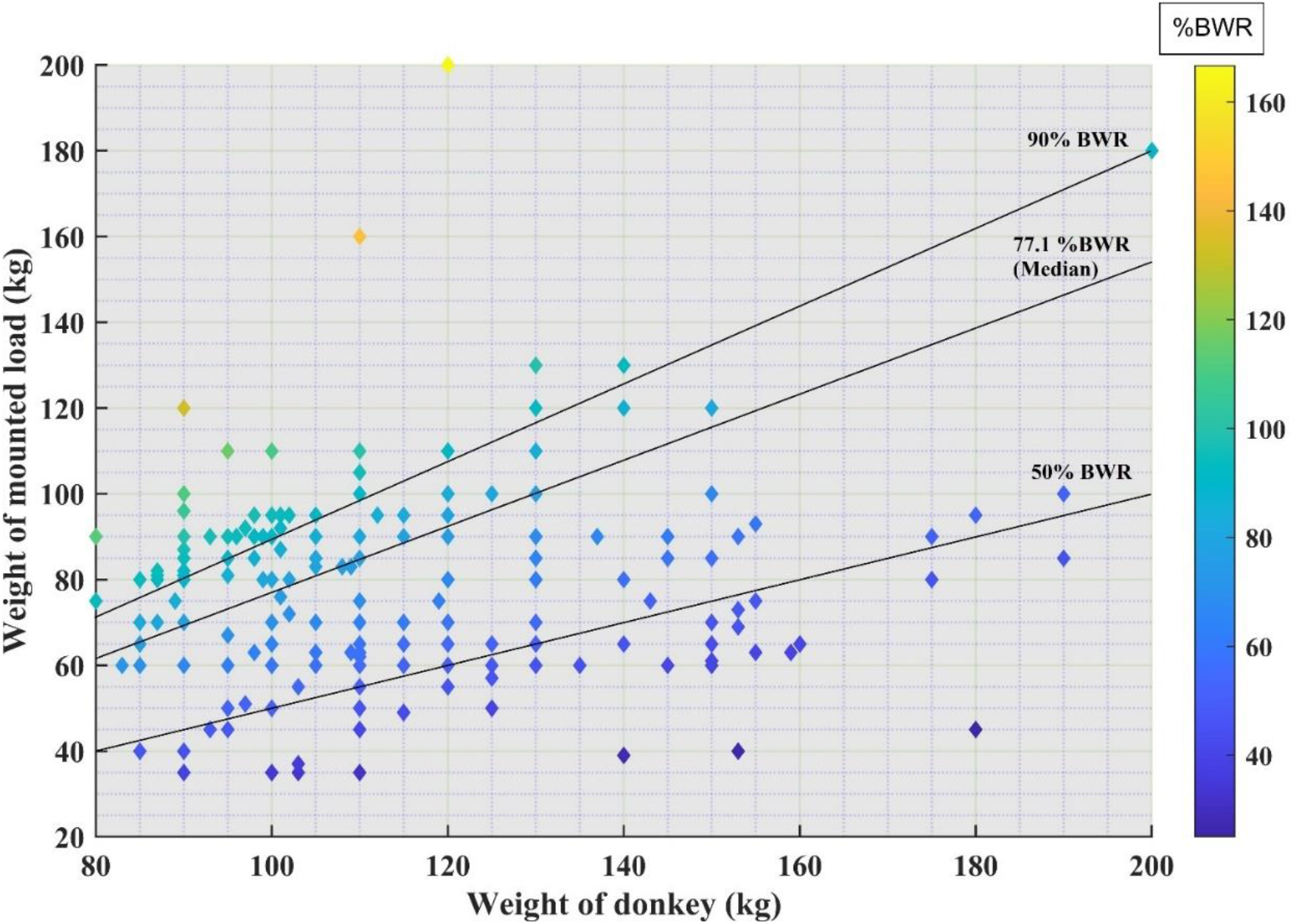
Donkey weight plotted against load carried. Lines represent 50% bodyweight ratio (BWR) carried, the median %BWR and high BWR (the upper quartile for %BWR). Lighter colors represent a higher %BWR.

### 3.3 Donkey owners and load carrying

Owners reported 44.0% (n=146) of donkeys were used for carrying construction-related material, 38.3% (n=127) were used for carrying agricultural-related material and 17.7% (n=59) were used for domestic goods. We found that 37.7% (n=125) of donkeys were working on flat terrain, 5.4% (n=18) on steep terrain, and 56.9% (n=189) on combined flat and steep terrain. Most donkeys (n=321, 96.7%) only walked during their routine daily work. In total, 41.6% (n=138) of the donkey owners reported that they routinely saw lameness in their donkeys while working (Table 2).

**Table 2:**
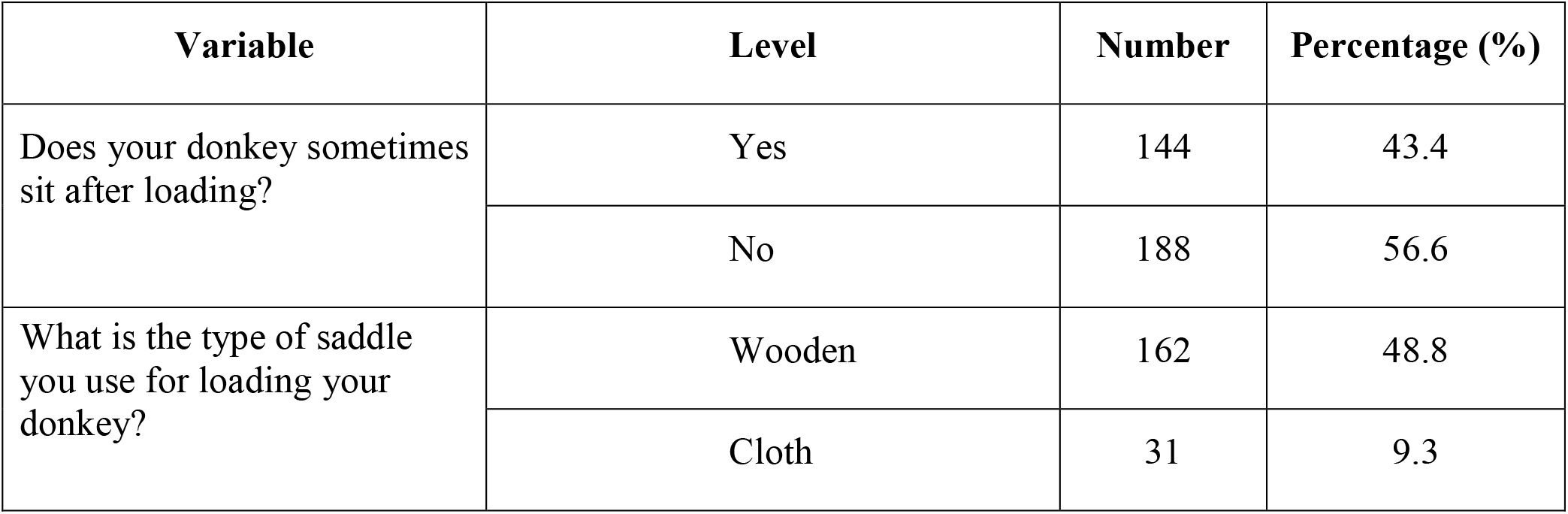
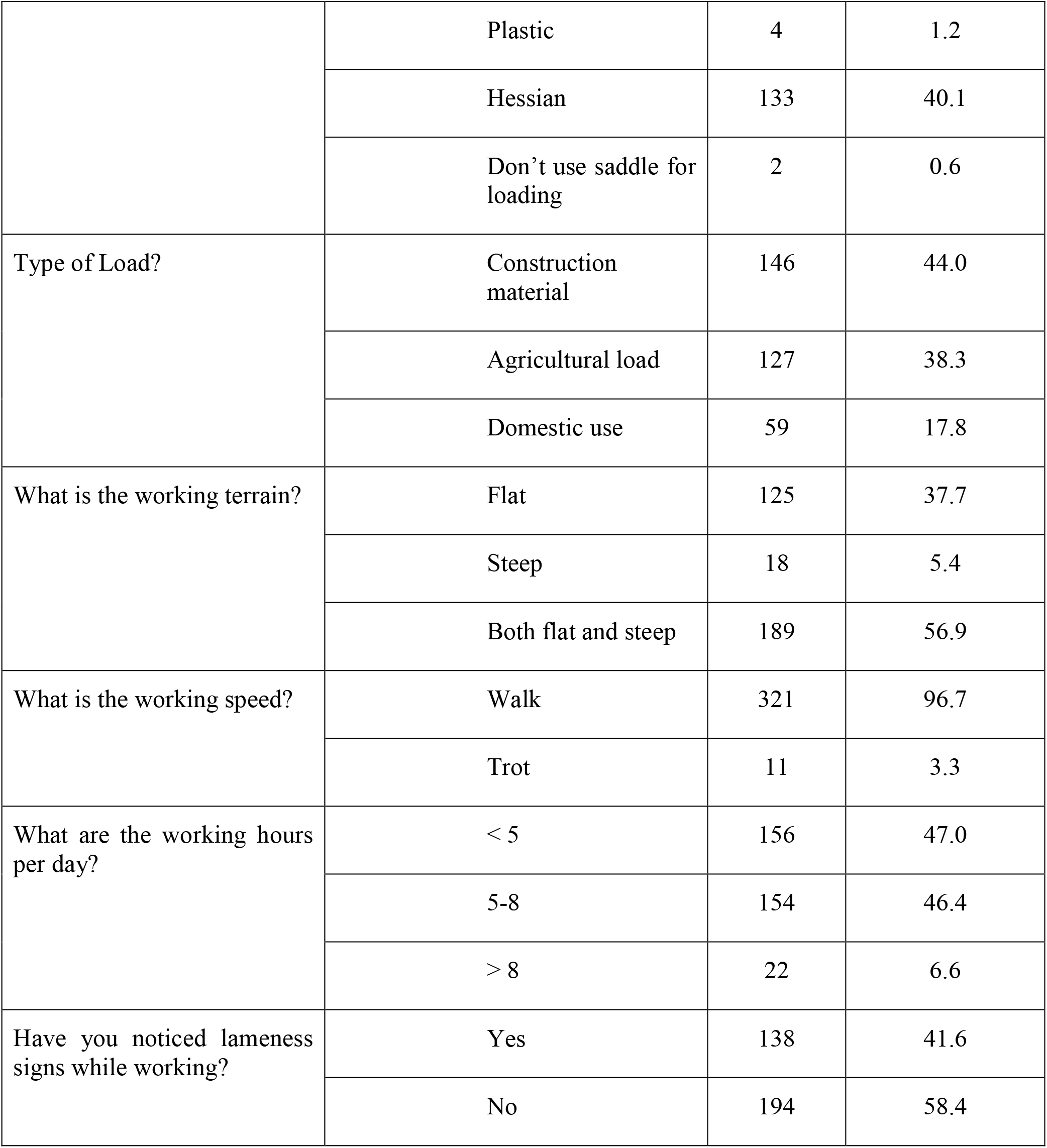
Practices of working donkey owners related to mounted load carrying

### 3.4 Multivariable regression modelling with respect to 50%, median and, high %BWR of load carrying donkeys

#### 3.4.1 Donkeys carrying 50 %BWR

Distance traveled (km), breed of donkeys, and sitting behavior after loading were all retained in the final model. Mixed breed donkeys were 2.57 (95% Confidence Interval (CI) 1.21 to 5.46) times more likely to carry loads of more than fifty percent of their bodyweight, compared to other breeds of donkeys (P = 0.01). Donkeys traveling over 8km per day were 7.16 (95% CI 1.47 to 34.79) times more likely to carry loads of more than 50% of their bodyweight, compared to donkeys traveling up to 3km per day (P = 0.01). Donkeys were 4.20 (95% CI 1.30 to 13.55) times more likely to sit when loaded if they were loaded with more than 50% of their bodyweight, compared with donkeys loaded with less weight (P = 0.01) (Table 3).

**Table 3:** Multivariable regression model with respect to 50% body weight ratio (%BWR) of load carrying donkeys

| Variable | Level | Donkey carrying below | Donkey carrying above | Odds ratio (OR) | 95% CI Lower | 95% CI upper | Wald P-value | Likelihood Ratio P-value |
| --- | --- | --- | --- | --- | --- | --- | --- | --- |

|  |  | **50**<br>**%BWR** | **50**<br>**%BWR** |  |  |  |  |  |
| --- | --- | --- | --- | --- | --- | --- | --- | --- |
| **Distance covered per day (Km)** | <4 | 24 | 89 | 1 |  |  |  | <0.001 |
|  | 4 to 8 | 16 | 38 | 0.50 | 0.23 | 1.08 | 0.08 |  |
|  | >8 | 2 | 163 | 7.16 | 1.47 | 34.79 | 0.01 |  |
| **Donkey Breed** | Other breeds | 24 | 60 | 1 |  |  |  | 0.01 |
|  | Mixed breed | 18 | 230 | 2.57 | 1.21 | 5.46 | 0.01 |  |
| **Does your donkey sometimes sit after loading?** | No | 38 | 150 | 1 |  |  |  | 0.008 |
|  | Yes | 4 | 140 | 4.20 | 1.30 | 13.55 | 0.01 |  |

#### 3.4.2 Donkeys carrying median %BWR

Area, age of donkey, type of load, earnings per day (PKR), and distance traveled (km) were all retained in the final model. The odds of carrying a load of more than the median %BWR in the sampled population of donkeys was higher if the donkey was working in a peri-urban (OR 2.78; 95% CI [1.16-6.63]) or urban area (12.82 OR; 95% CI [3.68-44.71]), compared to a rural area (P = <0.001). Younger donkeys aged between 1 and 5 years carried more than median weight compared with donkeys aged 15 or older (11.38 OR; 95% CI [1.10-117.20]; P = 0.03). Donkeys were loaded with more than the median weight if they carried construction materials (OR 5.41; 95% CI [1.69-17.26]; P = 0.004), (Table 4).

**Table 4:** Multivariable regression model with respect to median percent body weight ratio (%BWR) of load carrying donkeys

| **Variable** | **Level** | **Donkey carrying below the median** | **Donkey carrying above the median** | **Odds ratio (OR)** | **95% CI Lower** | **95% CI upper** | **Wald P-value** | **Likelihood Ratio P-value** |
| --- | --- | --- | --- | --- | --- | --- | --- | --- |
| **Area** | Per-urban | 22 | 104 | 2.78 | 1.16 | 6.63 | 0.02 | <0.001 |
|  | Rural | 135 | 28 | 1 |  |  |  |  |
|  | Urban | 9 | 34 | 12.82 | 3.68 | 44.71 | 6.26 |  |
| **Donkey age (years)** | 1 to 5 | 11 | 27 | 11.38 | 1.10 | 117.20 | 0.04 | 0.03 |
|  | 6 to 10 | 67 | 126 | 5.25 | 0.61 | 45.12 | 0.13 |  |
|  | 11 to 15 | 47 | 12 | 1.82 | 0.18 | 18.08 | 0.61 |  |
|  | >15 | 41 | 1 | 1 |  |  |  |  |
| **Type of Load?** | Domestic | 51 | 8 | 1 |  |  |  | <0.001 |
|  | Agriculture | 91 | 36 | 3.03 | 0.88 | 10.34 | 0.08 |  |
|  | Construction | 24 | 122 | 5.41 | 1.69 | 17.26 | 0.004 |  |
| **Earnings per day (PKR)** | <700 | 55 | 111 | 1 |  |  |  | 0.005 |
|  | 700 to 900 | 54 | 39 | 0.42 | 0.17 | 1.06 | 0.06 |  |
|  | >900 | 59 | 14 | 0.18 | 0.06 | 0.53 | 0.002 |  |
| **Distance covered per day (Km)** | <4 | 102 | 11 | 1 |  |  |  | <0.001 |
|  | 4 to 8 | 43 | 11 | 2.15 | 0.70 | 6.57 | 0.18 |  |
|  | >8 | 21 | 144 | 14.81 | 4.99 | 43.95 | 1.18 |  |

#### 3.4.3 Donkeys carrying high %BWR (90% BWR)

Area, age of donkey, breed, working terrain, and working hours per day were all retained in the final model. The odds of carrying a load of more than 90% of bodyweight was higher if the donkey was working in a peri-urban (OR 14.51; 95% (CI) [4.10-51.37]) or urban area (OR 8.38; 95% CI [2.15-32.67]), compared to rural areas (P ≤0.001). Mixed breed donkeys were 17.92 (95% CI 2.40 to 133.87) times more likely to carry loads of more than 90 percent of their bodyweight, compared to other breeds of donkeys (P = 0.005). Donkeys working for more than 8 hours a day were 26.31 (95% CI 4.11 to 168.52) times the odds of carrying load of 90% of BWR, compared to donkeys working less than 5 hours per day (Table 5).

**Table 5:**
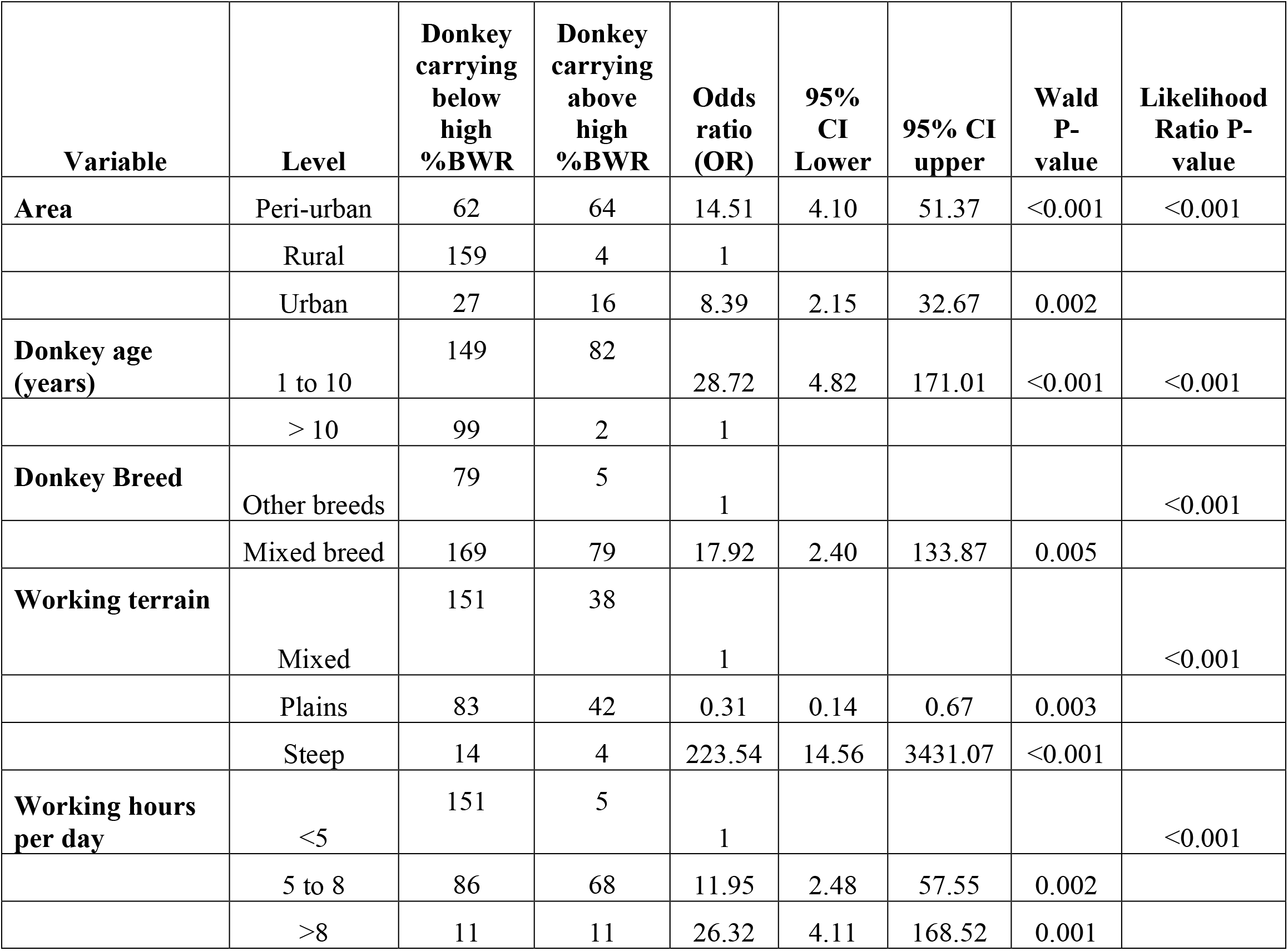
Multivariable regression model with respect to high percent body weight ratio (%BWR) of load carrying donkeys

## 4 Discussion

Studying load carrying in working donkeys is important because high workload and unsafe practices contribute to poor working donkey welfare (3,5). We explored mounted load carrying by donkeys in Pakistan and the factors associated with the weight of load carried. Despite overloading being an important donkey welfare problem (5), the current report is the first to elucidate factors associated with mounted loads carried under field conditions. One quarter of donkeys carried loads equal to 90% or more of their own bodyweight, with some donkeys estimated by their owners to be carrying more than 150% of their bodyweight. Factors including the type of load carried, the breed and age of the donkeys, and their location of work were associated with how much load the donkey carried. The variables associated with the loading of donkeys in this population have previously been associated with the poor welfare of working donkeys more broadly (5,11,24–28).

Overloading based on current recommendations (50% BWR) (15) was common, with the majority (87.4%) of donkeys reported to carry more than 50% BWR. The weight of donkeys in our study was comparable to a previous report of draught donkeys in Pakistan (17), suggesting that the data are robust. The weight of mounted loads found in our research is also similar to previous investigations from Ethiopia (11) and India (29). However, it has also recently been suggested that donkeys should not carry more than one third of their bodyweight (11). Further, experimental research has suggested that donkeys can travel further, for longer and with less physiological impact if they are loaded with 40% to 50% of their bodyweight (15). Moreover, guidelines for donkeys working on beaches in the United Kingdom mandate carrying not more than 28% of bodyweight (16). Compared to horses, donkeys are currently carrying much higher mounted loads as the maximum permissible load-carrying limits suggested for native Japanese horses is 29% (30,31), for Yonagunai ponies is 33% (32), and for Taishuh ponies is 43% of their bodyweight (31). Furthermore, a rider should not weigh more than 10% of the horse’s bodyweight in the UK, but in the US, this limit is doubled to 20% of the horse’s weight (16).

In urban areas, donkey owners were more likely to load their donkeys to more than 75% and more than 90% of their bodyweight compared to rural areas. The area a working donkey lives in is a known factor for poor working equine welfare (7,25,26), as rural donkeys usually had fewer lesions on their body than urban donkeys and a larger proportion of urban donkeys showed moderate to severe gait deviation (i.e., lameness) than rural donkeys (7,33). Moreover, rural donkeys work less than those in urban areas (4). Unfortunately, these authors did not define “work less” in terms of a lighter loaded weight, shorter working hours or distance traveled. However, this finding may be why fewer welfare concerns were raised for rural donkeys (7). Furthermore, donkeys in rural and urban settings have different roles within these communities and face different welfare challenges (4,7). Due to the differences in practices between urban and rural areas, determining the welfare and socio-economic value of working donkeys in different parts of the same territory is crucial.

The type of load carried (construction, agricultural, or domestic), was associated with the weight of mounted load. Donkeys working for the transportation of construction-associated load carry more weight than donkeys carrying agricultural loads. There is currently no research comparing the type of load carried and the weight of that load. However, working donkeys that transport different types of loads experience different impacts on their health and welfare (7,24,25,27,34–37). For example, donkeys used in brick transport are 2.5 times more likely to have moderate to deep skin lesions and 3.4 times more likely to have sole surface abnormalities than those used for other purposes (25). We hypothesise that because brick is a dense material, more bricks will fit on the back of a donkey than other materials, leading to heavier loads being carried when compared to less dense agricultural or domestic loads.

Most donkeys worked for less than eight hours and covered a median distance of 8km (ranges, 1-30km) per day. The daily working hours of donkeys varies in Ethiopia (20), Mexico (24), Egypt (27), and Nepal (37). In our study, donkeys working for a greater number of hours and covering more distance per day carried more weight. This is associated with the type of workload; donkeys working for transportation of construction associated load usually carry more weight, work for longer hours, and cover more distance than donkeys working for domestic or agricultural work, as they typically carry less, for a shorter period of time and over a shorter distance. Donkeys transporting agricultural load work less than donkeys involved with other types of work (4,25). Moreover, donkeys work for up to twelve hours a day in Ethiopia (11), and cover a distance of more than 30km a day in Morocco (26). The duration of work and distance travelled increases, it compromises donkey welfare (11,26,27). Longer working hours and distance to be covered in addition to high mounted load are likely to lead to fatigue, and fatigue from overworking and overloading can compromise donkey welfare and productivity (27).

Donkey age was associated with load carried in all models when donkeys were carrying more than 50% of their bodyweight. Younger animals between 1 and 5 years of age carried more load compared to older animals. In the UK it is recommended that donkeys be at least four years old before starting work (12). Donkeys may appear mature at the age of two, but they are not skeletally mature until they are three to four years old, and it has been suggested that donkeys should not carry weight or work until they were five to six years old to avoid osteoarthritic changes due to overworking (38,39). Previous research has found gait abnormalities, hoof abnormalities, tendon, joint swelling, and other load-associated injuries are more prevalent in older working donkeys (11,20,25); we suggest that this is because they have been carrying higher levels of mounted loads throughout their young lives, and they face multiple complex issues in their older age.

In our survey, 42% of donkey owners reported seeing lameness while their donkey was working. However, a more in-depth lameness examination by a veterinarian or other appropriately trained professional would be needed to confirm this. In comparison, visual signs of lameness were observed in 15% of working equids by experts in Mexico (24), while gait abnormalities in working equids reported by experts in a wide range of countries range from 17.1% to 99.2% (25). A recent study of working donkeys pulling carts in the Faisalabad region of Pakistan found that 96% of donkeys were lame when examined by a veterinarian, despite examination being conducted while the donkey was still in harness (40). Owners in our survey reported less lameness previous reports from Pakistan; this could be due to differences in areas within Pakistan or may be due to donkey owner abilities to identify lameness. This assumption is based on surveys of horses, which have repeatedly demonstrated that owners report a lower prevalence of lameness and gait asymmetry than experts (41,42). However, donkey owners have suggested work overload as a potential cause for lameness in Ethiopia (43) and Pakistan (40), and mule owners also recognize this issue (20). However, lameness is one of the main welfare issues reported in working equids globally (5,7,17,25,35,40,43,44) and this is an area for important future targeted owner education.

Donkey owners reported that donkeys carrying more than 50% of their bodyweight were more likely to sit after loading irrespective to type of load and area of work. However, in experimental research, donkey sitting behavior needs further investigation to evaluate its validity as a load quantifying factor in working donkeys. This study’s two major limitations are convenience sampling and owner-reported weights, both of which are unavoidable in this context. Because the data are based on answers gathered during owner interviews, the accuracy of the data must be carefully considered (26). Furthermore, the reliability of ‘owner information’ has not been validated and may be imperfect or biased due to owner reporting of perceived ‘correct’ answers (26).

This is a starting point for the development of evidence-based recommendations for donkey loading. It is clear based on the use and role of the donkeys in this study that recommending donkeys are only loaded at 50% of bodyweight will be a practical failure, if there is a lack of understanding of the motivations and perceptions of owners around donkey loading and the socio-economic role that load carrying donkeys play for donkey owners. While the welfare of the donkey is important, and a consideration here, donkey welfare can only be improved alongside community recognition of the issue, and a general improvement of human living conditions.

## 5 Conclusion

Our research has provided valuable information on the demographics of working donkeys, and the factors associated with mounted load carried by working donkeys in Pakistan. Factors including type of load carried, the breed, sitting behavior, and age of the donkeys, and their location of work were associated with how much load the donkey carried. Overloading based on current recommendations (50% BWR) was common, and 87.4% of donkeys were carrying more than 50% of their bodyweight in the survey region. As overloading is one of the most common welfare issues in working donkeys, this is an area in which future education efforts should be targeted.

## 7 Author Contributions

All authors were involved in the preparation of the manuscript and gave final approval of this manuscript. All authors have read and agreed to the published version of the manuscript.

## 9 Acknowledgments

We thank the local equine veterinarians, without whom this study would not have been possible, and all the donkey owners participated in this study.

## 10 Supplementary Material

**S1 Table. Univariable regression model with respect to 50 percent body weight ratio (50 %BWR)**. Exposure variables with a likelihood ratio test (LRT) P-value <0.20 were selected for inclusion in the multivariable model. (DOCX)

**S2 Table. Univariable regression model with respect to median percent body weight ratio (median %BWR)**. Exposure variables with a likelihood ratio test (LRT) P-value <0.20 were selected for inclusion in the multivariable model. (DOCX)

**S2 Table. Univariable regression model with respect to 90 percent body weight ratio (high %BWR)**. Exposure variables with a likelihood ratio test (LRT) P-value <0.20 were selected for inclusion in the multivariable model (DOCX)

## 11 Data Availability Statement

The dataset generated for this study is available from the corresponding author upon request.

