## Supplementary Tables for "Welfare concerns for mounted load carrying by working donkeys in Pakistan"

### Supplementary Material

**Table S1: Univariable regression model with respect to 50 percent body weight ratio (50 %BWR).**

| Variable | Level | Odds ratio | 95% CI Lower | 95% CI upper | Wald P-value | Likelihood Ratio P-value |
|---|---|---|---|---|---|---|
| **Breed of donkey** | Indian | 0.16 | 0.03 | 0.68 | 0.01 | <0.001 |
| | Mixed breed | 1 | | | | |
| | Shinghari | 0.19 | 0.09 | 0.43 | 4.51 | |
| | Sperki | 0.21 | 0.08 | 0.57 | 0.002 | |
| | Cons | 12.78 | 7.91 | 20.64 | 2.25 | |
| **Breed of donkey** | Other breeds | 1 | | | | <0.001 |
| | Mixed breed | 5.11 | 2.60 | 10.03 | 2.09 | |
| | Cons | 2.5 | 1.55 | 4.01 | <0.001 | |
| **Does your donkey sometimes sit after loading?** | No | 1 | | | | <0.001 |
| | Yes | 8.86 | 3.08 | 25.48 | 5.09 | |
| | Cons | 3.94 | 2.76 | 5.63 | 4.02 | |
| **Type of Saddle** | Hessian | 3.74 | 1.48 | 9.44 | 0.005 | 0.02 |
| | Cloth | 1 | | | | |
| | Plastic | 1.43 | 0.13 | 15.51 | 0.77 | |
| | Wooden | 4.34 | 1.74 | 10.82 | 0.001 | |
| | Cons | 2.1 | 0.99 | 4.46 | 0.05 | |
| **Type of Load?** | Agriculture | 0.16 | 0.05 | 0.48 | 0.001 | <0.001 |
| | Construction | 1 | | | | |
| | Domestic | 0.06 | 0.02 | 0.18 | 1.04 | |
| | Cons | 35.5 | 13.14 | 95.89 | 1.91 | |
| **Working terrain** | Mix | 1 | | | | <0.001 |
| | Plain | 8.92 | 2.67 | 29.73 | <0.001 | |
| | Steep | 0.57 | 0.19 | 1.70 | 0.31 | |
| | Cons | 4.55 | 3.14 | 6.60 | 1.14 | |
| **working speed** | Walk | 1 | | | | 0.04 |
| | Trot | 4.25 | 1.19 | 15.22 | 0.02 | |
| | Cons | 1.75 | 0.51 | 5.98 | 0.37 | |
| **Working hours per day** | < 5 | 1 | | | | <0.001 |
| | 5 to 8 | 5.85 | 2.50 | 13.66 | 4.45 | |
| | > 8 | 5.85 | 0.76 | 45.09 | 0.09 | |
| | Cons | 3.59 | 2.45 | 5.25 | 4.45 | |

| | | | | | | |
|---|---|---|---|---|---|---|
| **Lameness while working** | No | 1 | | | | <0.001 |
| | Yes | 6.27 | 2.39 | 16.40 | <0.001 | |
| | Cons | 4.24 | 2.96 | 6.07 | 2.61 | |
| **Earnings per day (PKR)** | <480 | 0.69 | 0.26 | 1.87 | 0.47 | 0.64 |
| | 480 to 690 | 1 | | | | |
| | 700 to 900 | 0.78 | 0.28 | 2.12 | 0.62 | |
| | >900 | 0.53 | 0.19 | 1.43 | 0.21 | |
| | Cons | 9.57 | 4.39 | 20.85 | 1.31 | |
| **Distance covered per day (Km)** | <4 | 1.56 | 0.74 | 3.26 | 0.24 | <0.001 |
| | 4 to 8 | 1 | | | | |
| | 9 t0 17 | 17.68 | 3.87 | 80.78 | <0.001 | |
| | >17 | 1 | | | | |
| | Cons | 2.37 | 1.32 | 4.26 | 0.004 | |
| **Distance travelled (km)** | <4 | 1.56 | 0.74 | 3.26 | 0.23 | <0.001 |
| | 4 to 8 | 1 | | | | |
| | >8 | 34.31 | 7.56 | 155.61 | 4.57 | |
| | Cons | 2.37 | 1.32 | 4.26 | 0.004 | |
| **Area** | Rural | 6.38 | 2.41 | 16.84 | <0.001 | <0.001 |
| | Peri-urban | 1 | | | | |
| | Urban | 3.51 | 1.02 | 12.05 | 0.04 | |
| | Cons | 3.79 | 2.60 | 5.53 | 4.61 | |
| **Donkey age (years)** | 1 to 5 | 13.13 | 1.60 | 107.42 | 0.02 | <0.001 |
| | 6 to 10 | 3.67 | 1.57 | 8.58 | 0.002 | |
| | 11 to 15 | 1.25 | 0.50 | 3.16 | 0.63 | |
| | >15 | 1 | | | | |
| | Cons | 2.82 | 1.41 | 5.60 | 0.003 | |
| **Donkey sex** | Female | 1 | | | | 0.5 |
| | Male | 1.25 | 0.65 | 2.38 | 0.50 | |
| | Cons | 6.14 | 3.87 | 9.74 | 1.22 | |

**Table S2: Univariable regression model with respect to median percent body weight ratio (median %BWR).**

| **Variable** | **Level** | **Odds ratio** | **95% CI Lower** | **95% CI upper** | **Wald P-value** | **Likelihood Ratio P-value** |
|---|---|---|---|---|---|---|
| **Breed of donkey** | Indian | 0.33 | 0.079 | 1.33 | 0.12 | <0.001 |
| | Mixed breed | 1 | | | | |
| | Shinghari | 0.19 | 0.09 | 0.38 | 5.42 | |
| | Sperki | 0.05 | 0.01 | 0.23 | 9.84 | |
| | Cons | 1.53 | 1.18 | 1.97 | 0.001 | |

| | | | | | | |
|---|---|---|---|---|---|---|
| **Breed of donkey** | Other breeds | 1 | | | | <0.001 |
| | Mixed breed | 6.50 | 3.56 | 11.86 | 1.04 | |
| | Cons | 0.23 | 0.13 | 0.40 | 1.93 | |
| **Does your donkey sometimes sit after loading?** | No | 1 | | | | <0.001 |
| | Yes | 9.94 | 5.94 | 16.61 | 1.93 | |
| | Cons | 0.38 | 0.27 | 0.52 | 3.72 | |
| **Type of Saddle** | Hessian | 7.17 | 2.87 | 17.95 | 2.53 | <0.001 |
| | Cloth | 1 | | | | |
| | Plastic | 1.14 | 0.10 | 12.78 | 0.91 | |
| | Wooden | 2.48 | 1.01 | 6.09 | 0.05 | |
| | Cons | 0.29 | 0.12 | 0.67 | 0.004 | |
| **Type of Load?** | Agriculture | 0.08 | 0.04 | 0.14 | 9.79 | <0.001 |
| | Construction | 1 | | | | |
| | Domestic | 0.03 | 0.01 | 0.07 | 3.08 | |
| | Cons | 5.08 | 3.28 | 7.87 | 3.31 | |
| **Working terrain** | Mix | 1 | | | | <0.001 |
| | Plain | 8.19 | 4.80 | 13.96 | 1.06 | |
| | Steep | 0.58 | 0.18 | 1.85 | 0.36 | |
| | Cons | 0.49 | 0.36 | 0.66 | 3.69 | |
| **Working speed?** | Walk | 1 | | | | 0.76 |
| | Trot | 1.21 | 0.36 | 4.03 | 0.76 | |
| | Cons | 0.83 | 0.25 | 2.73 | 0.76 | |
| **Working hours per day** | < 5 | 1 | | | | <0.001 |
| | 5 to 8 | 28.65 | 15.51 | 52.88 | 7.63 | |
| | > 8 | 20.71 | 6.93 | 61.86 | 5.71 | |
| | Cons | 0.16 | 0.10 | 0.25 | 4.02 | |
| **Lameness while working** | No | 1 | | | | <0.001 |
| | Yes | 6.93 | 4.23 | 11.38 | 1.69 | |
| | Cons | 0.46 | 0.34 | 0.62 | 4.65 | |
| **Earnings per day (PKR)** | <480 | 1.63 | 0.84 | 3.14 | 0.14 | <0.001 |
| | 480 to 690 | 1 | | | | |
| | 700 to 900 | 0.44 | 0.23 | 0.82 | 0.01 | |
| | >900 | 0.14 | 0.07 | 0.30 | 4.05 | |
| | Cons | 1.64 | 1.03 | 2.63 | 0.04 | |
| **Distance covered per day (Km)** | <4 | 0.42 | 0.17 | 1.04 | 0.06 | <0.001 |
| | 4 to 8 | 1 | | | | |
| | 9 t0 17 | 18.50 | 7.79 | 43.95 | 3.86 | |
| | >17 | 47.56 | 16.42 | 137.78 | 1.11 | |
| | Cons | 0.25 | 0.13 | 0.50 | 5.46 | |
| **Distance traveled (km)** | <4 | 0.42 | 0.17 | 1.04 | 0.07 | <0.001 |

| | | | | | | |
|---|---|---|---|---|---|---|
| | 4 to 8 | 1 | | | | |
| | >8 | 26.80 | 11.98 | 59.96 | 1.18 | |
| | Cons | 0.25 | 0.13 | 0.49 | 5.46 | |
| **Area** | Rural | 22.79 | 12.33 | 42.12 | 1.92 | <0.001 |
| | Peri-urban | 1 | | | | |
| | Urban | 18.21 | 7.86 | 42.19 | 1.27 | |
| | Cons | 0.21 | 0.14 | 0.31 | 3.58 | |
| **Donkey age (years)** | 1 to 5 | 100.63 | 12.27 | 825.07 | 1.74 | <0.001 |
| | 6 to 10 | 77.10 | 10.37 | 573.01 | 2.18 | |
| | 11 to 15 | 10.47 | 1.30 | 84.00 | 0.03 | |
| | >15 | 1 | | | | |
| | Cons | 0.02 | 0.003 | 0.18 | <0.001 | |
| **Donkey sex** | Female | 1 | | | | <0.001 |
| | Male | 3.51 | 2.23 | 5.53 | 5.93 | |
| | Cons | 0.5 | 0.35 | 0.70 | 6.28 | |

**Table S3: Univariable regression model with respect to 90 percent body weight ratio (high %BWR).**

| **Variable** | **Level** | **Odds ratio** | **95% CI Lower** | **95% CI upper** | **Wald P-value** | **Likelihood Ratio P-value** |
|---|---|---|---|---|---|---|
| **Breed of donkey** | Mixed breed | 1 | | | | <0.001 |
| | Shinghari | 0.14 | 0.04 | 0.46 | 0.001 | |
| | Sperki | 0.19 | 0.04 | 0.77 | 0.02 | |
| | Cons | 0.47 | 0.36 | 0.61 | 2.41 | |
| **Breed of donkey** | Other breeds | 1 | | | | <0.001 |
| | Mixed breed | 7.38 | 2.88 | 18.95 | 3.21 | |
| | Cons | 0.06 | 0.02 | 0.15 | 2.16 | |
| **Does your donkey sometimes sit after loading?** | No | 1 | | | | <0.001 |
| | Yes | 2.57 | 1.55 | 4.28 | <0.001 | |
| | Cons | 0.21 | 0.14 | 0.31 | 7.11 | |
| **Type of Saddle** | Hessian | 6.93 | 1.58 | 30.38 | 0.01 | 0.01 |
| | Cloth | 1 | | | | |
| | Plastic | 4.83 | 0.33 | 70.40 | 0.25 | |
| | Wooden | 4.44 | 1.01 | 19.48 | 0.05 | |
| | Cons | 0.07 | 0.01 | 0.29 | <0.001 | |

| | | | | | | |
|---|---|---|---|---|---|---|
| **Type of Load?** | Agriculture | 0.44 | 0.25 | 0.76 | 0.003 | <0.001 |
| | Construction | 1 | | | | |
| | Domestic | 0.12 | 0.04 | 0.36 | <0.001 | |
| | Cons | 0.59 | 0.42 | 0.82 | 0.002 | |
| **Working terrain** | Mix | 1 | | | | 0.03 |
| | Plain | 2.01 | 1.20 | 3.36 | 0.01 | |
| | Steep | 1.13 | 0.35 | 3.64 | 0.83 | |
| | Cons | 0.25 | 0.17 | 0.36 | 2.91 | |
| **Working hours per day** | < 5 | 1 | | | | <0.001 |
| | 5 to 8 | 23.88 | 9.27 | 61.50 | 4.91 | |
| | > 8 | 30.2 | 8.90 | 102.45 | 4.56 | |
| | Cons | 0.03 | 0.01 | 0.08 | 6.53 | |
| **Lameness while working** | No | 1 | | | | <0.001 |
| | Yes | 2.86 | 1.71 | 4.76 | 5.32 | |
| | Cons | 0.20 | 0.14 | 0.30 | 1.09 | |
| **Earnings per day (PKR)** | <480 | 1.93 | 0.96 | 3.89 | 0.06 | 0.002 |
| | 480 to 690 | 1 | | | | |
| | 700 to 900 | 1.56 | 0.77 | 3.17 | 0.29 | |
| | >900 | 0.45 | 0.18 | 1.12 | 0.08 | |
| | Cons | 0.27 | 0.16 | 0.48 | 5.11 | |
| **Distance traveled (km)** | <4 | 0.34 | 0.07 | 1.58 | 0.17 | <0.001 |
| | 4 to 8 | 1 | | | | |
| | 9 t0 17 | 13.09 | 4.35 | 39.45 | 4.83 | |
| | >17 | 8.97 | 2.95 | 27.27 | <0.001 | |
| | Cons | 0.08 | 0.03 | 0.22 | 1.17 | |
| **Distance covered per day (Km)** | <4 | 0.34 | 0.07 | 1.58 | 0.17 | <0.001 |
| | 4 to 8 | 1 | | | | |
| | >8 | 10.94 | 3.77 | 31.67 | 1.05 | |
| | Cons | 0.08 | 0.03 | 0.22 | 1.17 | |
| **Area** | Rural | 41.03 | 14.33 | 117.48 | 4.49 | <0.001 |
| | Peri-urban | 1 | | | | |
| | Urban | 23.55 | 7.32 | 75.83 | 1.18 | |
| | Cons | 0.02 | 0.01 | 0.06 | 3.48 | |
| **Donkey age (years)** | 1 to 5 | 29.82 | 3.70 | 240.02 | 0.001 | <0.001 |

| | | | | | | |
|---|---|---|---|---|---|---|
| | 6 to 10 | 21.31 | 2.86 | 158.37 | 0.002 | |
| | 11 to 15 | 0.71 | 0.04 | 11.63 | 0.81 | |
| | >15 | 1 | | | | |
| | Cons | 0.02 | 0.01 | 0.17 | <0.001 | |
| **Donkey sex** | Female | 1 | | | | 0.62 |
| | Male | 1.13 | 0.69 | 1.87 | 0.62 | |
| | Cons | 0.31 | 0.21 | 0.46 | 1.66 | |
| **Working speed?** | Walk | 1 | | | | 0.57 |
| | Trot | 1.54 | 0.33 | 7.29 | 0.58 | |
| | Cons | 0.22 | 0.05 | 1.03 | 0.05 | |
